# Human Serum Albumin as a Hidden Source of Variability in IVF Culture Media

**DOI:** 10.1101/2025.05.29.656106

**Authors:** Markéta Nezvedová, Volodymyr Porokh, Tami Bočková, Václav Pustka, Drahomíra Kyjovská, Barbora Maierová, Soňa Kloudová, Pavel Otevřel, Zuzana Holubcová

**Affiliations:** RECETOX, Faculty of Science, Masaryk University, 625 00 Brno, Czech Republic; Department of Histology and Embryology, Faculty of Medicine, Masaryk University, 625 00 Brno, Czech Republic; Reprofit International – Clinic of Reproductive Medicine and Gynecology, 603 00 Brno, Czech Republic; Proteomics Core Facility, Central European Institute for Technology, Masaryk University, 625 00 Brno, Czech Republic

**Author notes:** Corresponding author, address: | Masaryk University, Faculty of Medicine, Department of Histology and Embryology, Masaryk University Campus, Kamenice 3, building F01B1, 62500 Brno, Czech Republic.

**Keywords:** IVF media composition, human serum albumin, spent culture medium, embryo culture, proteomics, protein biomarkers

## Abstract

Soluble proteins present in spent culture media (SCM) have been proposed as potential indicators of embryo viability. However, results across studies remain inconsistent, and the identity of reliable protein biomarkers is still unclear. To evaluate the potential of secretome profiling for non-invasive embryo assessment, we applied state-of-the-art mass spectrometry to analyze the protein composition of SCM from individual successful and unsuccessful human embryo cultures, alongside corresponding controls cultured under identical conditions. Surprisingly, both untargeted (nanoLC-TIMS-TOF-MS/MS) and targeted (UHPLC-QqQ-MS/MS) proteomic analyses consistently detected multiple human proteins in unconditioned media, indicating that these signals are not of embryonic origin. Undeclared proteins were also identified in media sampled before their first use, with considerable variability observed across production batches. Supplementation experiments demonstrated that this variability originates from the addition of plasma-derived human serum albumin, which introduces a range of contaminating proteins. In contrast, recombinant albumin did not contribute detectable serum proteins. These findings indicate that the protein background in commercial IVF media is an overlooked variable, not only compromising the reproducibility and interpretation of secretome analyses in research, but also contributing to unstandardized culture conditions in clinical IVF practice. While this study evaluated 13 production lots of monophasic media from two different manufacturers, the results underscore a broader need for transparency and standardization in IVF media composition. The adoption of fully chemically defined IVF media, free from purified serum components, would support more consistent embryo culture conditions and advance the discovery of biomarkers in SCM.

## INTRODUCTION

Embryo selection for transfer is a crucial step in the in vitro fertilization (IVF) process, directly influencing implantation success and treatment outcomes. Improving embryo selection has the potential to shorten time to pregnancy and reduce emotional and financial burdens for patients. Despite its importance, embryo selection remains challenging, as universally reliable and objective criteria for assessing embryo quality have yet to be established.

Morphological assessment remains the standard approach for embryo grading in clinical practice ^1–3^. While straightforward, this method is inherently subjective and provides limited predictive accuracy for implantation outcomes. Time-lapse imaging allows continuous monitoring of embryo developmental dynamics and assessment of morphokinetic parameters ^4^. However, a consensus has yet to be reached on the definition of normal developmental trajectories and the optimal timing of key embryonic events ^5–7^.

Preimplantation genetic testing (PGT), which involves the biopsy of embryonic cells to analyze chromosomal composition or genetic abnormalities, offers a higher degree of selection accuracy than morphology assessment alone. However, clinical utility of PGT for aneuploidy is debated ^8^. Biopsied cells may not fully represent the whole embryo’s genetic status due to mosaicism ^9,10^, and emerging evidence suggests that human embryos have the capacity to self-correct downstream from the blastocyst stage ^11^, potentially eliminating the benefit of PGT screening. In addition, genetic testing requires embryo cryopreservation to allow time for analysis and remains a labor-intensive, time-consuming, and costly procedure.

In search of a non-invasive alternative, spent culture medium (SCM) analysis has emerged as a promising approach for embryo selection. During in vitro culture, the medium accumulates biochemical signals resulting from embryonic secretion and uptake, including low-molecular-weight metabolites, proteins, short non-coding RNAs, and extracellular vesicles ^12–15^. By analyzing these molecular signatures, researchers and clinicians could gain valuable insights into embryo viability, implantation potential, and overall developmental competence without directly interfering with the embryo.

Among the molecules detectable in SCM, soluble protein factors hold significant potential for in vitro diagnostics. A range of cytokines and growth factors, including IL-1α, IL-1β, IL-6, IL-8, IL-10, TNFR1, VEGF-A, PLGF, LIF, and GM-CSF, have been proposed as potential indicators of embryo viability and fitness ^14,16–18^. Additionally, the presence of factors involved in gene expression regulation (JARID2) ^19^, cellular energy metabolism, and oxidative stress (APOA1) ^20,21^, may further reflect the developmental competence of the embryo. The secretion of proteins with immunomodulatory functions (HRG, HLA-G) ^22–25^ or roles in cell adhesion (EMMPRIN, EpCAM)^26^ may also contribute to embryo–maternal crosstalk, supporting implantation and early pregnancy establishment.

Despite the growing list of proposed protein biomarkers in SCM, it is important to note that most available data originate from immunodetection-based experiments ^22–30^, warranting cautious interpretation due to methodological limitations. While immunoassays are highly sensitive, their specificity is often limited by the affinity and selectivity of the antibodies used, which increases the risk of false-positive and nonspecific results^31^. Only a small number of investigations have employed mass spectrometry (MS), a technique well suited for the comprehensive identification of multiple protein species in complex biological samples ^18,19,21,32,33^. Recent advances have enhanced its sensitivity, allowing the detection of low-abundance analytes from individual embryo cultures and expanding the potential for biomarker discovery ^34^. Yet, all published MS-based studies analysing SCM protein composition have applied only untargeted approaches without validating protein identities using purified protein or peptide standards ^19,21,32^. The scarcity of robust supporting evidence, combined with methodological shortcomings, raises concerns about the reliability of the reported protein biomarkers.

In this study, we employed a state-of-the-art MS-based approach to analyze SCM from individual embryo cultures to identify protein fingerprints associated with embryo developmental competence. For this purpose, we have employed both discovery-based untargeted and targeted MS methods to confirm selected protein identification using standard analytes. Surprisingly, our analysis revealed that not only SCM but also unused commercial IVF media, labeled as containing only human serum albumin (HSA), harbored a substantial number of undeclared proteins, with notable variability across different media batches. This finding led us to investigate the source of this protein background, which poses a significant challenge for proteomic analysis of the human embryo secretome.

## MATERIALS AND METHODS

### Embryo Culture and Sample Collection

A total of 53 spent culture media (SCM) samples from 22 IVF cycles were provided by a collaborating IVF unit for protein composition analysis. Following hormonal stimulation and oocyte retrieval, all mature oocytes were fertilized via intracytoplasmic sperm injection (ICSI), according to the clinic’s routine procedures. The inseminated oocytes were transferred to individual culture wells in an EmbryoSlide+ ic8 culture dish (#16454 Vitrolife, Sweden), prefilled with embryo culture medium (20 μL per well) and covered with prewarmed paraffin oil (Ovoil, #10029; Vitrolife, Sweden). This cultivation setup ensured that the medium in each well remained completely separated. The multiwell dishes were placed in a time-lapse incubator (EmbryoScope+, Vitrolife, Sweden) under stable environmental conditions at 37 °C, 5% O₂, and 6% CO₂. The same medium was maintained throughout the incubation period without replacement or renewal (Fig 1A). Embryo development was evaluated daily by experienced clinical embryologists as part of routine practice. SCM samples were categorized based on embryo development outcomes: “SPENT+”, representing media from successful cultures where embryos reached the blastocyst stage (on day 5 or day 6 post-ICSI), and “SPENT−“, representing media from unsuccessful cultures where embryos were arrested at the cleavage stage (Fig 1B). The corresponding control sample (CONTROL) consisted of media from a spare well in the same multiwell dish, which was incubated under identical conditions but without embryos. At the end of the six-day culture period (day 6), 10 μL of the embryo-free medium was carefully collected into low-protein-binding microcentrifuge tubes. Besides, 300 μl of unused medium was collected immediately after the first vial opening for reference (BLANK). All collected samples were frozen at -20 °C and transferred to the research laboratory, where they were stored at -80°C until analysis.

**Fig 1:**
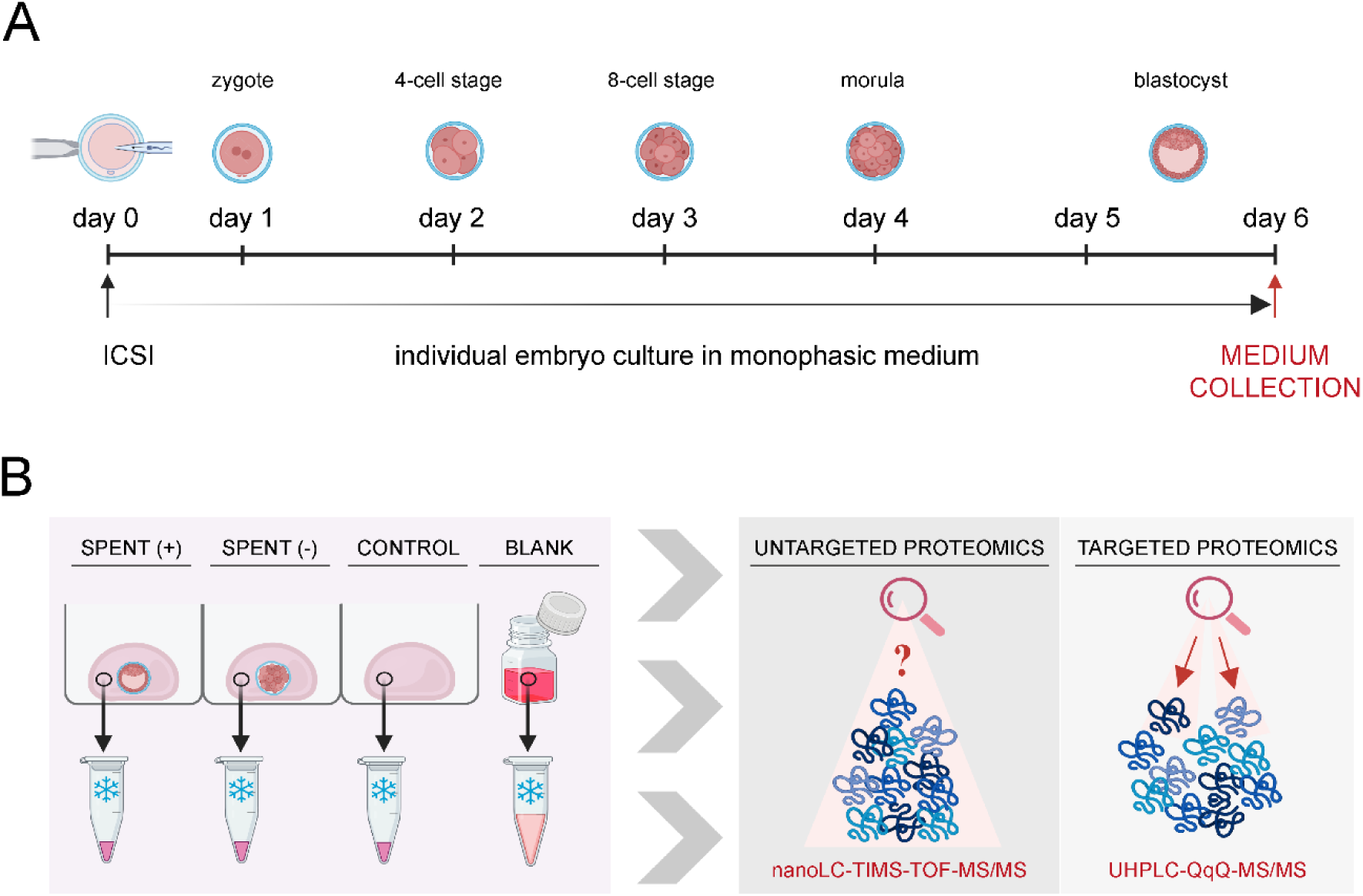
Workflow of embryo culture media analysis. (A) Timeline of embryo cultivation. At day 0, the intracytoplasmic sperm injection (ICSI) was performed, and embryos were then cultivated in a monophasic embryo culture medium. On day 6, spent media were collected for further analysis. (B) Strategy of embryo culture media analysis. Spent media were collected from individually cultured embryos whose development was assessed as good-quality (+) or arrested (-). Incubated media without embryo presence were collected as control samples (CONTROL), and non-incubated fresh media were collected as blank samples (BLANK). All four types of media samples were subjected to untargeted and targeted proteomic analyses using the corresponding system of liquid chromatography coupled with mass spectrometry.

### The embryo culture media and supplements

Two commercially available IVF culture media brands—Continuous Single Culture Complete (CSCM-C) (Cat. No. 90165, Irvine Scientific/FUJIFILM) and SAGE 1-Step™ (Cat. No. 67010010, CooperSurgical)—were analyzed in this study across 13 different production batches (**Supplementary Fig 1**). The media were used for human embryo cultivation as part of routine IVF practice within their declared shelf life. Storage and handling followed the manufacturer’s instructions and adhered to good laboratory practices.

To investigate potential sources of protein contamination in IVF media, we analyzed a protein-free IVF medium (Continuous Single Culture, CSCM, Cat. No. 90164, Batch: (10)0000031344; Irvine Scientific/Fujifilm) as a negative control, as well as media prepared by supplementing the same protein-free medium with either recHSA (Cellastim S™, Cat. No. A9731, Batch: 0000311138; Merck) or purified serum-derived HSA. The three different batches of serum-derived HSA included one product from Vitrolife (Cat. No. 10064: HSA-batch 1: 047982) and two product batches from Irvine Scientific/Fujifilm (Cat. No. 9988: HSA-batch 2: 0000021432 and HSA-batch 3: 0000021429).

### Untargeted Proteomics

#### Sample Preparation

Protein samples of SCM and controls (∼50 μg of total protein) were processed using the filter-aided sample preparation (FASP) method as previously described ^35^, with digestion performed using 1 μg of trypsin (proteomics grade, EMS0005, Merck). The resulting peptides were extracted into LC-MS vials using 2.5% formic acid (FA) in 50% acetonitrile (ACN) and 100% ACN with the addition of polyethylene glycol (final concentration 0.001%) ^36^ and were subsequently concentrated in a SpeedVac concentrator (Thermo Fisher Scientific).

Blank media samples (all media samples not exposed to embryos) were processed using in-solution digestion and selected over FASP to better assess the background protein composition and minimize contamination risk. This method involves fewer handling steps and is therefore less prone to introducing external protein contaminants. An aliquot of each sample (∼50 μg of total protein) was diluted using 50mM ammonium bicarbonate (ABC) buffer (pH 8) with n-dodecyl-beta-maltoside (DDM, final conc. 0.01%) and subjected to reduction with dithiothreitol (DTT, final conc. 5 mM) and alkylation with iodoacetamide (IAA, final conc. 10 mM). 1 μg of trypsin (proteomics grade, EMS0005, Merck) was used for the digestion (4 hours, 37 °C) of the protein mixture, and the obtained peptide solution was acidified with 5% FA (final concentration 0.5%).

#### Instrumentation and Data Acquisition

LC-MS/MS analyses of all peptides were performed using the UltiMate 3000 RSLCnano system (Thermo Fisher Scientific) coupled to the timsTOF Pro 2 mass spectrometer (Bruker). Prior to LC separation, tryptic digests were online concentrated and desalted using a trapping column (Acclaim PepMap 100 C18, 300 μm ID, 5 mm length, 5 μm particles, Thermo Fisher Scientific). The trap column was washed with 0.1% trifluoroacetic acid (TFA), and peptides were eluted in backflush mode onto an analytical column (Aurora C18, 75 μm ID, 250 mm length, 1.7 μm particles, Ion Opticks) using a 120-minute gradient program at a flow rate of 200 nL/min, increasing from 3% to 42% mobile phase B (0.1% FA in 80% ACN), followed by a system wash with 80% mobile phase B. Equilibration of both the trapping and analytical columns was performed prior to sample injection. The analytical column was installed in the Captive Spray ion source (Bruker) with the temperature set to 50°C according to the manufacturer’s instructions. The spray voltage and sheath gas were set to 1.4 kV and 1, respectively.

MS data were acquired in data-independent acquisition (DIA) mode, with a base method covering an m/z range of 100–1700 and 1/k₀ range of 0.6–1.4 V×s×cm⁻². The enclosed DIAparameters.txt file defined an m/z 400–1000 precursor range with equal window sizes of 21 Th, utilizing two steps for each parallel accumulation-serial fragmentation (PASEF) scan and a cycle time of 100 ms, locked to 100% duty cycle.

#### Data Processing

DIA data were processed using DIA-NN (version 1.8.1) ^37^ in library-free mode, searching against a modified cRAP database (based on http://www.thegpm.org/crap; 111 sequences in total) and the UniProtKB protein database for Homo sapiens (https://www.uniprot.org/proteomes/UP000005640; version 2023/11, containing 20,596 protein sequences). Carbamidomethylation was set as a fixed modification, while trypsin/P was specified as the enzyme with one missed cleavage allowed and a peptide length range of 7–30 amino acids. False discovery rate (FDR) control was set at 1%. MS1 and MS2 accuracies, as well as scan window parameters, were optimized based on median values from initial test searches. Match-between-runs (MBR) was enabled. The mass spectrometry proteomics data have been deposited to the ProteomeXchange Consortium via the PRIDE partner repository ^38^ with the dataset identifier PXD063245 (reviewer access token: bKwOy4H7oTKP).

### Targeted Proteomics

#### Sample Preparation

Total protein concentrations were determined using the bicinchoninic acid (BCA) protein assay. Collected spent media (SCM), control, and blank media samples were diluted fourfold with ammonium bicarbonate buffer (AmBic, ≥99.5% purity, 100 mM) to achieve a final protein concentration of approximately 1 μg/μL. The samples were vortexed for 10 seconds at 2000 rpm (VELP Scientifica). For blank samples, two to four aliquots (technical replicates) were prepared and processed individually, following the same procedure as for the SCM and control samples.

Aliquots of 40 μL (containing 40 μg of total protein) were reduced with 20mM 1,4-dithiothreitol (DTT, ≥99% purity) in 2.5mM AmBic for 10 minutes at 95°C. Alkylation was then performed using 40 mM iodoacetamide (≥99% purity) in 2.5 mM AmBic for 30 minutes in the dark at room temperature. Protein digestion was carried out using mass spectrometry-grade trypsin gold (Cat. # 5280; Promega) at a 1:40 ratio (enzyme: total protein content, w/w). The samples were sealed with Parafilm and incubated overnight (16 hours at 37 °C) with gentle shaking.

Before terminating the enzymatic reaction, protein digests were spiked with a mixture of isotopically labeled synthetic crude peptides (SpikeTides L, JPT Technologies, final concentration ∼12.8 nM). Digestion was halted by adding 200 μL of 2% formic acid (FA, ∼98% purity, MS grade). The samples were then purified using solid-phase extraction (Oasis PRiME HLB, 30 mg, Waters Corp). The washing phase was performed with 300 μL of 2% FA, followed by two elution phases with 200 μL of 50% acetonitrile (ACN, LC-MS grade) containing 2% FA. Finally, the samples were dried using a vacuum evaporator system (Genevac).

#### Instrumentation and Data Acquisition

Dried samples were reconstituted in 10 μL of 5% ACN with 0.1% FA, matching the initial conditions of the liquid chromatography (LC) gradient. Positive ion detection was used for the analysis of targeted peptides using an ultra-high-performance liquid chromatography-tandem mass spectrometry (UHPLC-MS/MS) system (1290 Infinity II, Agilent 6495B) in selected reaction monitoring (SRM) mode. A sample volume of 3 μL, corresponding to 12 μg of total protein, was injected into a C18 pre-column (CSH 1.7 μm, 2.1 mm × 5 mm, Waters Corp) and separated on a C18 analytical column (Peptide CSH 1.7 μm, 2.1 mm × 100 mm, Waters Corp). The mobile phases consisted of water with 0.1% FA (phase A) and 95% ACN with 0.1% FA (phase B), with a flow rate of 0.3 mL/min. A linear gradient elution was applied, starting at 5 % of phase B and increasing to 30 % over 25 minutes, followed by a rapid increase to 95% B from 25.5 to 30 minutes before returning to 5% B from 31 to 35 minutes. The ion source parameters were set to a temperature of 200 °C and a capillary voltage of 3500 V. The SRM protein assays and instrument-specific libraries for targeted protein analysis were created using publicly available protein databases (www.nextprot.org and www.srmatlas.org).

#### Data processing

Data were manually inspected and processed using the Skyline software (online available). Selected single quantifier transitions were used to calculate relative concentrations using the following formula: peak area of native peptide/peak area of corresponding standard peptide × final concentration of spiked standard peptide. Mass spectrometry data from the targeted proteomics experiment have been deposited in the Panorama Public repository and are accessible via the permanent link: https://panoramaweb.org/GHihM3.url. The dataset is also available through the ProteomeXchange Consortium with the identifier PXD063038 (https://proteomecentral.proteomexchange.org/cgi/GetDataset?ID=PXD063038), and is associated with the DOI: https://doi.org/10.6069/vkze-gj62. The reviewer account details: Email: <u>,</u> Password: FL!y?xOkG0IU+r.

## Data analysis and visualization tools

Protein MaxLFQ intensities reported in the DIA-NN main report were further processed using a reproducible software container environment (Omics Workflows; https://github.com/OmicsWorkflows, version 4.7.7a). The complete processing workflow is available upon request. Briefly, the pipeline included (a) removal of low-quality precursors and contaminant protein groups, (b) imputation of missing values, (c) calculation and log₂ transformation of protein group MaxLFQ intensities, and (d) differential expression analysis using the LIMMA statistical test. Proteins with an adjusted p-value < 0.05 and a fold change > 2 were considered significantly differentially abundant. To visualize the data from the untargeted analysis, we included only results on proteins identified with one or more unique peptides, while those identified solely through shared peptides were excluded. Detected proteins were categorized using selected Gene Ontology categories, and their normalized intensities were visualized in a barplot from the Matplotlib module (Matplotlib 3.7.2, Python 3.11.5)^39^. Principal component analysis (PCA) plots and protein concentration graphs based on targeted proteomic data (Fig 2C, **Supplementary Fig 2**, **Fig 3**, **Supplementary Fig 3**, **Fig 4**, **Supplementary Fig 4**, **Fig 5B, C**, and **Supplementary Fig 5**) and performed statistical tests (Fig 3**, Supplementary Fig 3**) were created using GraphPad Prism (version 10.4.1). Heatmaps were generated from normalized and scaled protein intensities using the R package ComplexHeatmap (version 2.22.0) in R version 4.4.2. The protein selection used for each heatmap was generated as a set passing the standard threshold of 0.05 with its adjusted (Benjamini-Hochberg method) p-value from a differential expression analysis of proteins across different batches or manufacturers respectively, using a linear modeling approach (Limma v3.62.2) with Empirical Bayes moderation applied to improve variance estimation. The adjacent clustering dendrogram uses the complete hierarchical clustering method based on Euclidean distances between the samples. Fig 1 (https://BioRender.com/79qxcoz), Fig 5A (https://BioRender.com/p5q7vfs), and Fig 6 (https://BioRender.com/wc1ndat) were created with BioRender and published as BioRender illustrations under the CC-BY 4.0 license for open access. Icons in **Supplementary Fig 1** were taken from Biorender (https://BioRender.com).

**Fig 2:**
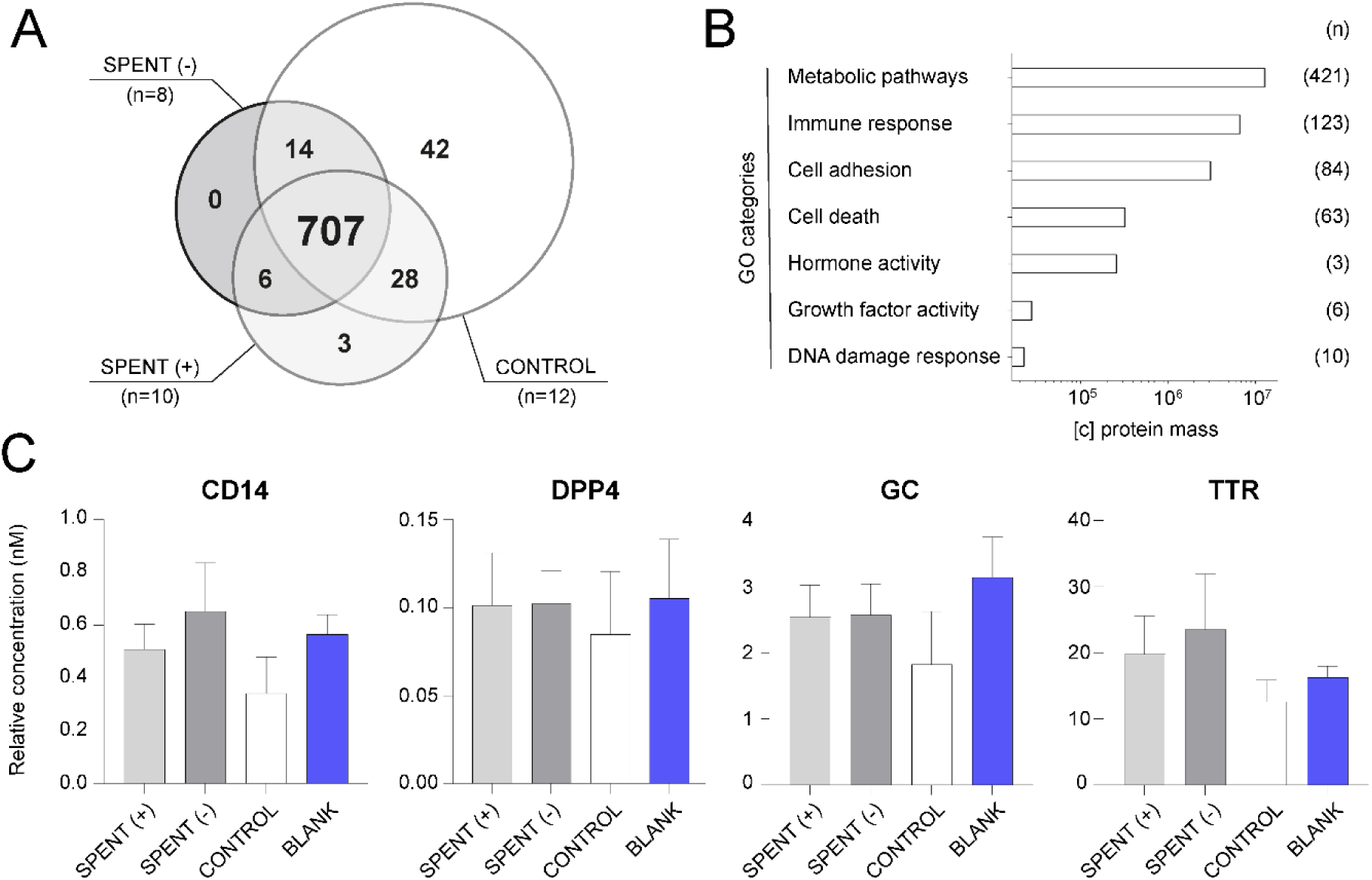
Protein analysis of embryo culture media. (A) Untargeted analysis of SCM samples from good-quality (SPENT+) and poor-quality (SPENT−) embryos, along with their corresponding controls (CONTROL), revealed that only a low number of protein markers were unique to each sample group, while the majority of proteins were shared across groups (central intersection); (n) indicates number of samples per group. (B) Gene Ontology (GO) categorization of proteins detected in control samples by untargeted proteomics. The x-axis represents the mean protein mass size, and (n) indicates the number of proteins assigned to each GO category. (C) Targeted analysis of four selected proteins (CD14, DPP4, GC, and TTR) in a subset of SCM samples (n=17) from good-quality (SPENT+, n=8) and poor-quality (SPENT−, n=9) embryo cultures, corresponding controls (CONTROL, n=5), and blank samples of unused media (BLANK, n=8 technical replicates). The results for an additional 10 proteins are presented in **Supplementary Fig2**.

**Fig 3.**
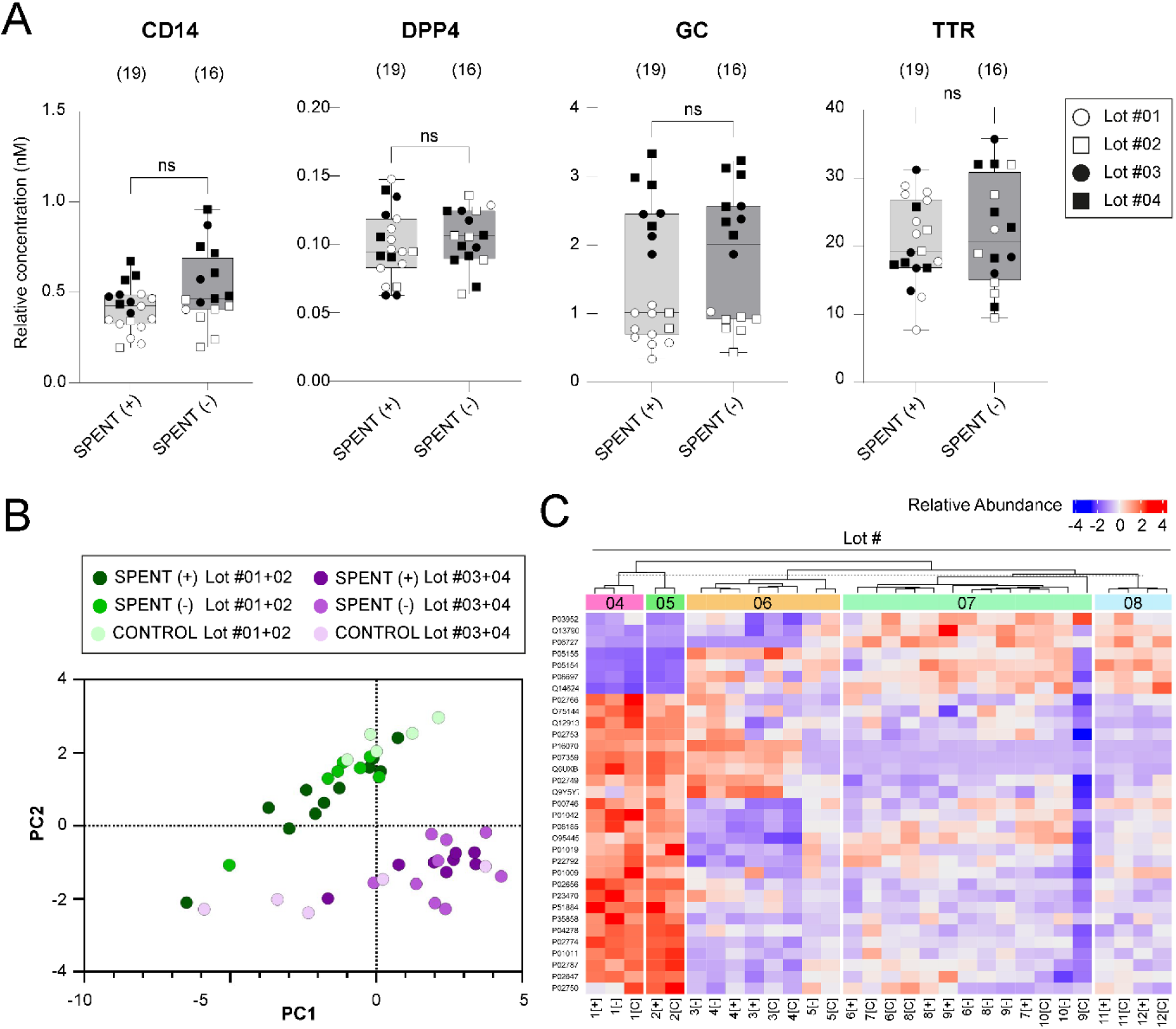
: Protein analysis of spent culture media with respect to the media Lot number. (A) Targeted analysis of four selected proteins (CD14, DPP4, GC, and TTR) in SCM from good-quality (SPENT+) and poor-quality (SPENT−) embryos, categorized based on the Lot number (Lot #01-04) of the culture media used. "ns" indicates statistically non-significant differences (Mann-Whitney test, p>0.05). (n) denotes the number of samples per group. The results for an additional 10 proteins are presented in **Supplementary Fig3**. (B) Principal component analysis (PCA) of 14 proteins identified in SCM from good-quality (SPENT+) and poor-quality (SPENT−) embryo cultures and corresponding control media, revealing two distinct groups associated with the Lot number of the culture media. (C) Heatmap showing relative abundance of proteins identified by untargeted proteomic analysis in SCM [SPENT (+) vs. SPENT (–)] and corresponding control media samples across five different production lots. Protein identifiers are listed on the left, and the color scale indicates z-score normalized abundance values. Samples are grouped by hierarchical clustering based on similarity in protein abundance profiles.

## RESULTS

### Types of samples and analysis workflow

To investigate the protein composition of spent culture media (SCM) and its potential role in assessing embryo development, we analyzed samples collected from 22 IVF cycles performed under controlled laboratory conditions (Fig 1A). Embryos were individually cultured in a monophasic embryo culture medium, ensuring each embryo remained in an isolated microenvironment. Of the 53 SCM samples collected from the collaborating IVF clinic, 29 were derived from successful cultures where embryos developed to the blastocyst stage (SPENT+), while 24 originated from unsuccessful cultures where embryos were arrested at the cleavage stage (SPENT−). The SCM samples, along with their corresponding control and blank samples, were subjected to state-of-the-art untargeted and targeted proteomic analysis using mass spectrometry technology (Fig 1B).

### Protein analysis revealed the presence of undeclared proteins in embryo culture media

Our pilot untargeted analysis of a subset of SCM samples (n=18) detected a substantial number of protein species present in both successful (SPENT+, n=10) and unsuccessful (SPENT-, n=8) culture samples. However, we did not identify any proteins that could reliably distinguish between the two groups (Fig 2A**, Supplementary Table 1**). Unexpectedly, the vast majority of detected proteins were also found in control samples (n=12) cultured in the absence of embryos, indicating that these proteins are not embryo-derived. Interestingly, a few proteins were exclusively detected in control samples, indicating possible degradation of these proteins in the presence of embryos.

While HSA is the only protein officially declared in commercial IVF media, our analysis revealed the presence of additional, undeclared proteins. This finding prompted further investigation into their origin and potential effects on embryo development. Based on Gene Ontology (GO) categories, we found that many proteins detected in control samples are implicated in essential biological processes, including cellular metabolism, immune response, and cell adhesion, and could therefore influence embryo development and implantation potential. Notably, some of these proteins possess hormonal or growth factor activity, further suggesting a potential impact on embryo viability (Fig 2B**, Supplementary Table 2**).

Building on our findings from the untargeted analysis, we further investigated the presence of undeclared proteins in SCM from (un)successful embryo cultures and corresponding controls using a targeted proteomics approach. From the proteins identified in the untargeted analysis, we selected 14 candidates involved in cellular metabolism, immune response, cell adhesion, and hormone transport (**Supplementary Table 3**). Our analysis confirmed the presence of all candidate proteins in SCM and corresponding controls. No consistent trend was observed between SPENT+ and SPENT− samples, as some proteins showed higher levels in the SPENT+ group, while others were more abundant in the SPENT− group. Importantly, all proteins of interest were also detected in blank media (Fig 2C, **Supplementary** Figure 2), confirming that they are not embryo-derived but instead originate from the unused culture medium itself. These findings align with our untargeted analysis data, reinforcing that the presence of proteins in SCM does not necessarily reflect embryonic secretion.

### Variations in Protein Concentrations Are Influenced by Media Lot

Next, we normalized the targeted analysis data from SCM samples to their corresponding controls to assess differences between successful (SPENT+) and unsuccessful (SPENT-) embryo cultures. No significant differences in protein fold-changes relative to control media were detected between the two groups (data not shown). Instead, we observed relative concentration variations in specific analytes, namely, CD14, GC, APOA1, BCHE, and IGHG2, that were associated with the culture media batch used, here referred to as Lot #01–#04 (Fig 3A, **Supplementary Fig 3**). Principal component analysis (PCA) further supported these findings; the PCA plot revealed two distinct clusters, separating samples based on the media batch (Lot #01 + Lot #02 versus Lot #03 + Lot #04), irrespective of the embryo developmental success (Fig 3B). This separation suggests that the observed differences in protein levels are driven by batch-specific variations in the commercially produced culture media rather than by embryo-derived signals.

To investigate this effect further, we revisited the untargeted proteomic data from individual SCM samples (Fig 2A**, Supplementary Table 1**), focusing on variability associated with different media lots. We observed separation driven by differential protein abundance profiles and identified 33 protein species that contributed most prominently to the overall variability. As illustrated in the heatmap (Fig 3C), hierarchical clustering reveals distinct grouping patterns that closely align with specific media lots, regardless of embryo culture outcome. Notably, six of the proteins showing media lot-associated variation (APOA1, APOH, RBP4, TTR, GC, and CD44) were also quantified in the targeted protein panel, reinforcing the consistency of this batch effect across analytical platforms.

### Blank Media Protein Variations Across Producers and Batches

Following these findings, we extended our analysis to evaluate blank media samples collected at the collaborating clinic over time. We applied our targeted protein detection methodology to quantify preselected proteins in these samples, which were taken after the first opening of a new package. A total of nine samples, representing different batches from two manufacturers, were analyzed (Product A: Lot 1-4; Product B: Lot 6-9). Our results demonstrate that levels of undeclared proteins vary between manufacturers and among lots from the same product (Fig 4A, **Supplementary Fig 4**).

**Fig 4:**
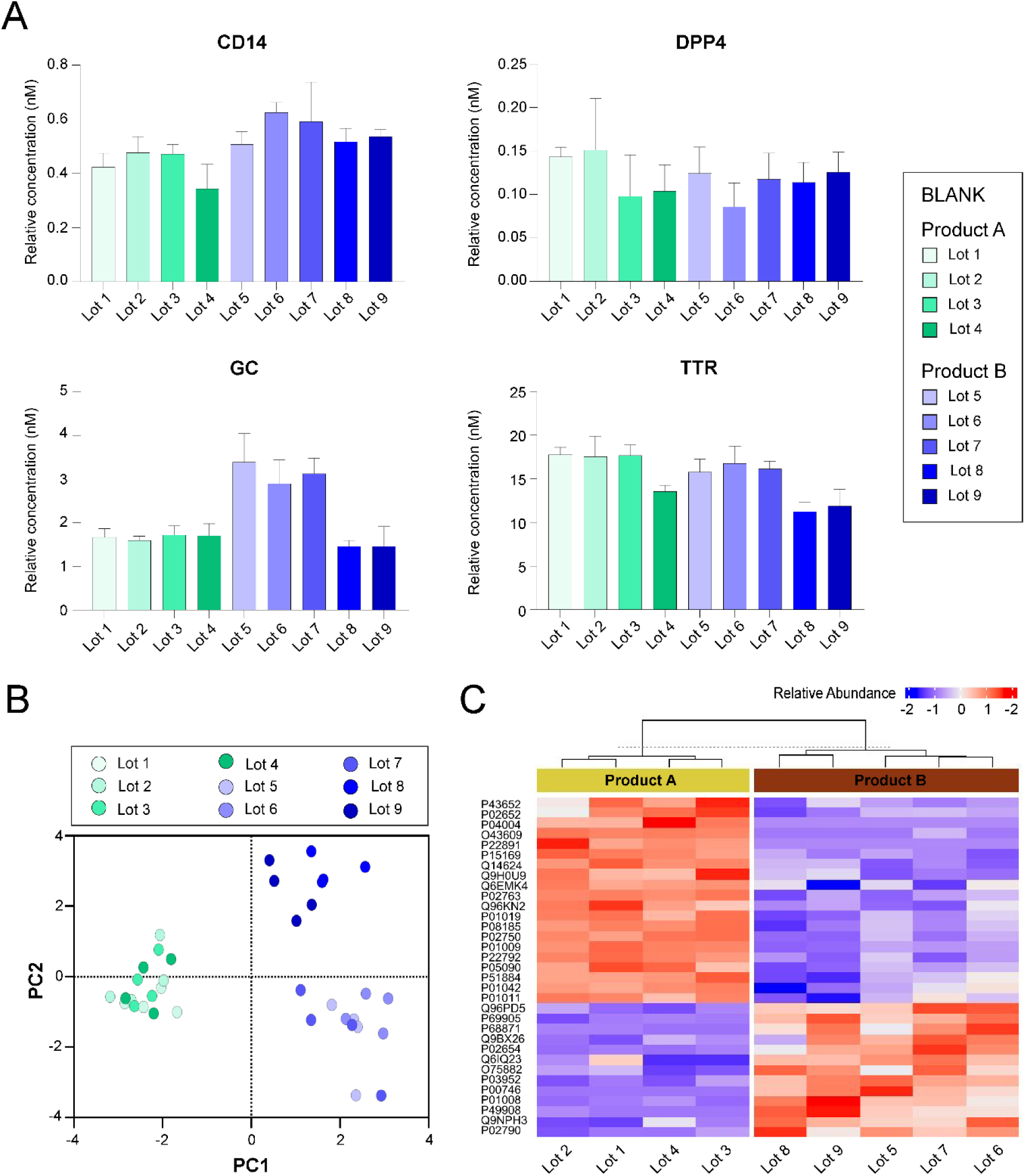
Protein analysis of blank media samples. (A) Levels of four selected proteins (CD14, DPP4, GC, and TTR) detected by targeted analysis in 9 Lots of embryo culture media from two different producers—Product A (shades of green, n=4 technical replicates per Lot) and Product B (shades of blue, n=4 technical replicates per Lot). The results for an additional 10 proteins are presented in **Supplementary Fig4**. (B) Principal component analysis (PCA) of 14 protein levels across different blank media Lots, highlighting variability between products. (C) Heatmap showing relative abundance of proteins identified by untargeted proteomic analysis in blank media samples from nine different lots (Product A: Lot 1–4; Product B: Lot 5–9). Protein identifiers are listed on the left, and the color scale indicates z-score normalized abundance values. Samples are grouped by hierarchical clustering based on similarity in protein abundance profiles.

**Fig 5:**
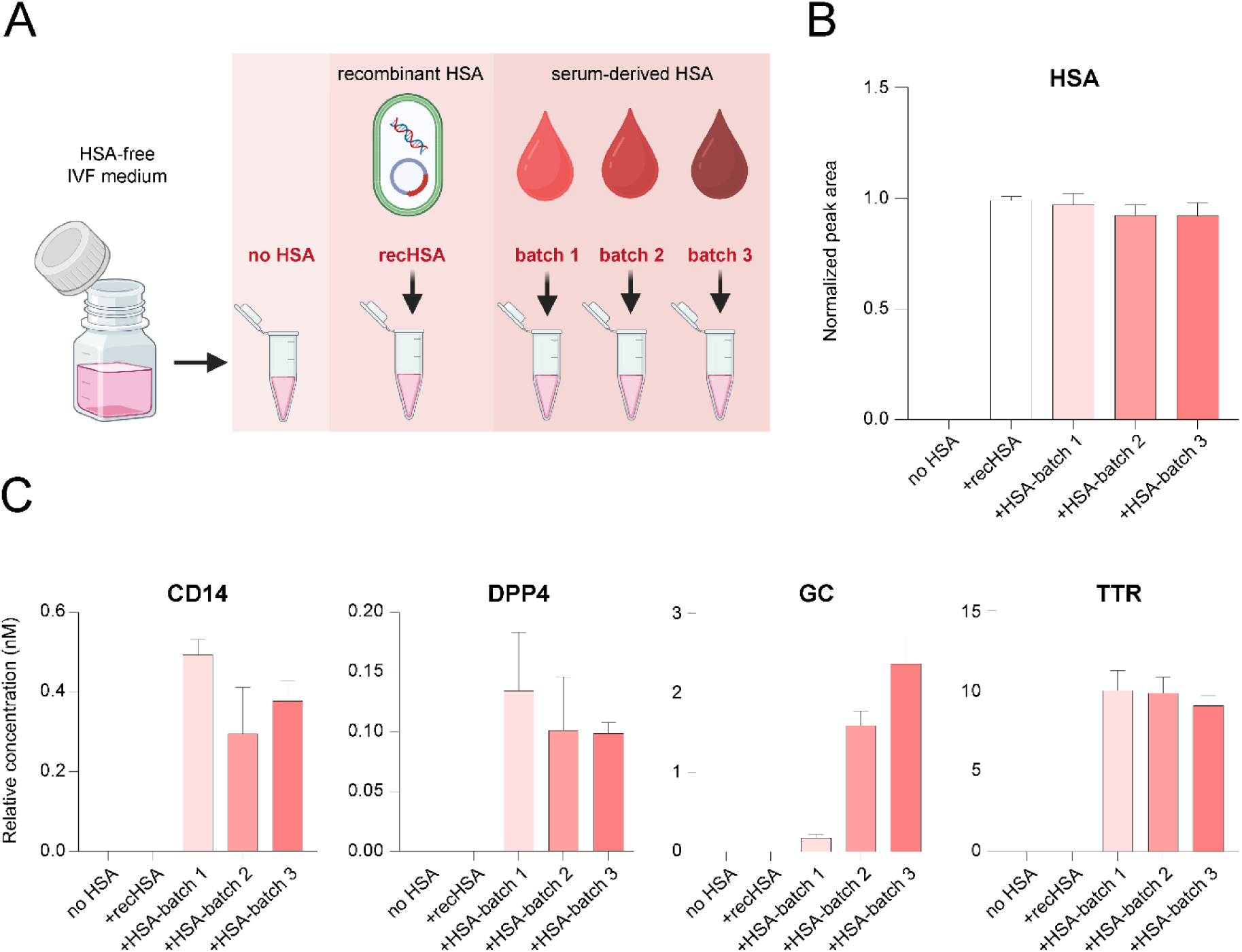
Investigation of protein background sources in embryo culture media. (A) Experimental design illustrating the analysis of HSA-free IVF medium (no HSA), recombinant HSA (recHSA), and serum-derived HSA from three different production batches (batch 1, batch 2, batch 3) added to HSA-free media; n=4 technical replicates per sample type. (B) Levels of HSA detected in the analyzed samples, showing its presence in all samples supplemented with either recHSA or serum-derived HSA, and its absence in the HSA-free medium (no HSA). (C) Targeted analysis of four selected proteins (CD14, DPP4, GC, and TTR) in all types of samples. These proteins were detectable only in samples supplemented with serum-derived HSA, with varying levels depending on the HSA production batch, but were absent in samples containing recHSA or no HSA. The results for the remaining analyzed proteins are shown in **Supplementary** Figure 5.

**Fig 6:**
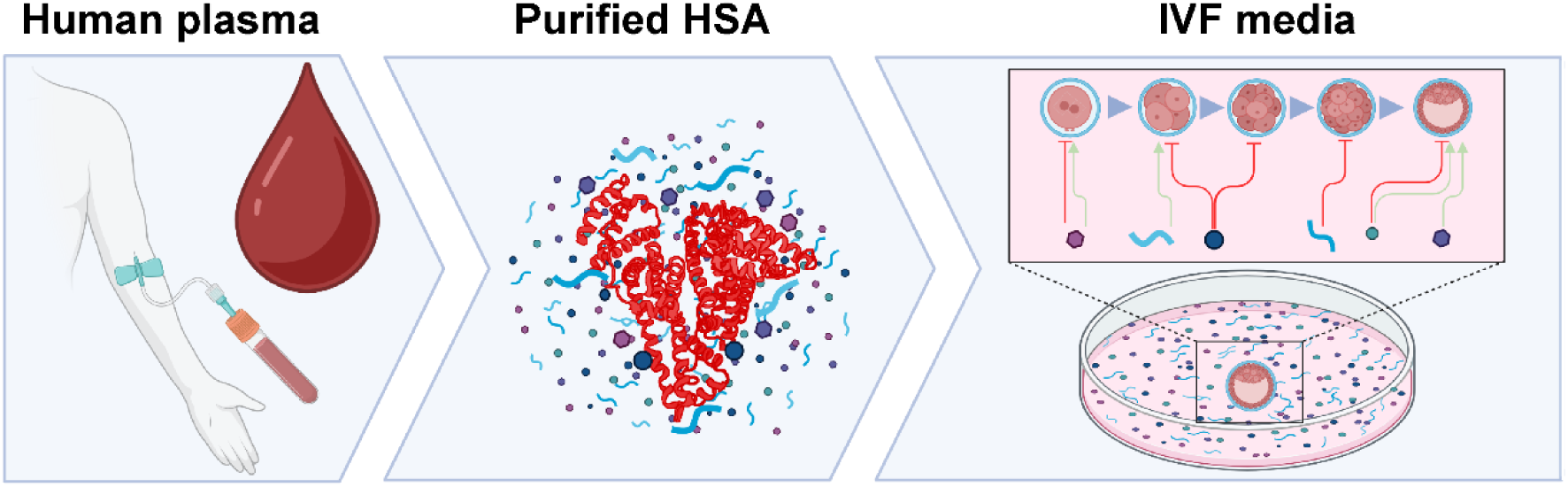
Schematic representation of HSA as a source of protein contamination in IVF media. Human serum albumin (HSA) derived from human plasma may carry residual contaminants, which are introduced into IVF media and can impact embryo culture and the proteomic profile of spent culture media.

PCA of the targeted analysis of 14 proteins identified three distinct clusters, confirming differences in protein levels between producers (green and blue shades). Additionally, PCA revealed two clusters corresponding to variations among batches from the same producer (Fig 4B).

To complement the targeted analysis, we performed untargeted proteomic profiling on the same set of blank media samples. This analysis revealed distinct proteomic signatures associated with each manufacturer, with Product A and Product B forming clearly separated clusters (Fig 4C). Numerous proteins differed consistently between the two product groups, reflecting manufacturer-specific composition. Variability was also observed within each product, with Product B showing greater batch-to-batch differences. These findings support the targeted analysis and reinforce the conclusion that both inter-manufacturer and intra-manufacturer variability exist in blank media protein content, even before embryo culture.

### Blood-Derived Albumin as the Source of Undeclared Proteins in Commercial IVF Media

HSA is the only protein officially declared to be present in the IVF culture medium. We hypothesized that the purification process used to obtain serum HSA from human blood may not yield a completely pure protein. Consequently, supplementing media with serum-derived HSA may introduce traces of other plasma proteins that remain associated with albumin throughout the fractionation and purification process.

To test this hypothesis, we supplemented a protein-free medium with HSA from different sources. The experimental conditions included: (1) unsupplemented protein-free medium, (2) medium supplemented with recombinant HSA (recHSA), and (3) medium supplemented with serum-derived HSA purified from human blood. The serum-derived HSA was obtained from two different manufacturers, with one supplier providing two distinct batches. This resulted in three separate preparations of blood-derived HSA added to protein free media (Fig 5A). A control experiment confirmed the presence of HSA in all supplemented samples, while it was absent in the unsupplemented condition (Fig 5B).

Targeted proteomic analysis demonstrated that the selected contaminating proteins were detected exclusively in samples supplemented with blood-derived HSA. These proteins were completely absent in both the unsupplemented medium and the condition supplemented with recombinant HSA, indicating that recHSA did not introduce any of the measured contaminants (Fig 5C**, Supplementary Fig5**). Furthermore, the levels of these proteins varied between the three blood-derived HSA preparations, suggesting batch-specific differences in contaminant load. Untargeted proteomic analysis confirmed these observations and identified additional serum-associated proteins introduced by HSA supplementation. While all blood-derived HSA preparations were found to contain a core set of shared proteins, each batch also included distinct proteins not present in the others, indicating inter-lot variability in the contaminant profile (**Supplementary Fig6**).

Together, these results support our hypothesis that undeclared proteins are introduced into the culture media through the use of blood-derived HSA (Fig 6).

## DISCUSSION

Our study uncovered a previously underappreciated complexity and variability in the protein composition of commercial IVF culture media before embryo exposure. Using a comprehensive proteomic approach, we identified numerous undeclared proteins in unconditioned culture media, with significant compositional variability across production lots and manufacturers. Further, by supplementing protein-free media with HSA from various sources, we demonstrated that serum-derived HSA is a major contributor to this unintended protein contamination. These findings highlight a critical yet often overlooked source of non-standardization in embryo culture conditions. A protein background complicates the interpretation of proteomic signals in SCM and hinders the identification of reliable embryo-derived biomarkers.

Although human embryos have been shown to develop in protein-free media ^40^, most IVF cultures include protein supplements, which are believed to support cell growth by stabilizing pH, maintaining osmotic balance, providing antioxidant protection, and facilitating cell adhesion, metabolism, and signalling. The protein composition of embryo culture media has evolved significantly over the past few decades (reviewed in ^41^. Early IVF attempts relied on protein-rich physiological fluids such as human tubal fluid, which mimicked the in vivo environment but posed challenges due to variability, limited availability, and biosafety concerns. The shift to blood-derived albumin offered a more stable and biochemically defined protein source. Over time, HSA purified from human serum became the standard protein supplement in embryo culture media.

Most ready-to-use IVF media declare HSA (5 mg/mL) as the sole protein additive. This leads to the common assumption that any additional proteins detected in SCM must be embryo-derived. However, our analysis detected over 700 protein species also in control media samples cultured without embryos, indicating that many proteins in SCM do not originate from the embryo. This result is in line with earlier work by Dyrlund and colleagues ^42^, who reported only over 100 contaminating proteins in IVF media. The lower number in their study likely reflects stricter identification criteria, requiring confirmation by three or more peptides, whereas our approach includes all proteins detected with at least one unique proteotypic peptide to capture the full spectrum of potentially present species.

Importantly, we observed that the protein composition of unconditioned IVF media varies depending on the HSA batch used for supplementation, leading to inconsistent culture conditions. This undermines the distinction between embryo-secreted proteins and media-derived protein background, raising concerns about the validity of earlier protein studies that lacked proper media-only controls. For instance, while we detected proposed biomarkers such as HRG and APOA1 in SCM, these were also present in control samples, suggesting a media-related origin. Moreover, APOA1 levels varied significantly across blank media batches, further supporting the idea that its abundance reflects manufacturing differences rather than embryo biology.

The discrepancies between our MS-based proteomics data and existing immunoassay studies are not unexpected, given the fundamental differences in detection methods. Immunoassays are prone to cross-reactivity, underscoring the importance of including appropriate controls ^31^. The divergence from previous MS-based studies may be due to their reliance on untargeted analysis without targeted validation. In contrast, we complemented untargeted analysis with a targeted proteomic approach, incorporating four technical replicates to robustly confirm the presence of 14 selected protein candidates in this complex biological matrix.

Another important consideration is the high sensitivity of modern proteomic techniques, which is advantageous for detecting low-abundance proteins but also makes them susceptible to identifying trace contaminants introduced during sample processing. Any proteomic study may, therefore, detect proteins that are not originally present in the sample but are introduced through handling, reagents, or equipment. In particular, media manipulation during dish preparation and SCM sampling in a clinical laboratory can represent potential points of entry for external proteins. This highlights the need for well-controlled experimental design and standardized sample processing protocols in proteomic studies to avoid mistaking contamination for true biological signals.

To pinpoint the source of background proteins, we demonstrated that supplementation with serum-derived HSA introduces a range of contaminating proteins, likely originating from serum. Albumin is the most abundant blood protein known for its capacity to bind and transport other molecules, including amino acids, small lipids, vitamins, hormones, and drugs. Current purification methods, typically involving plasma fractionation, do not guarantee the complete removal of HSA-associated molecules ^43^. As a result, incompletely purified HSA may introduce a batch-specific mix of co-purified serum proteins into embryo culture media.

The biological impact of molecular factors “hitchhiking” on HSA remains unclear, warranting further investigation into their potential role in embryo development. Several undeclared proteins detected in our control media were classified as having hormonal or growth factor functions, suggesting they could actively modulate developmental processes. Identifying HSA-associated embryotrophic factors could support the refinement of IVF media composition by preserving beneficial components and eliminating harmful or unnecessary contaminants. In a mouse model, HSA-bound fatty acids were shown to influence embryonic growth, indicating that albumin-associated compounds can actively shape developmental outcomes ^44^. Moreover, clinical studies reported a correlation between the type of IVF media and perinatal outcomes ^45,46^, suggesting that media composition can have lasting effects beyond the preimplantation period. Recent analysis has revealed substantial differences in the composition of commercially available media, pointing to a lack of standardization among producers ^47^. In this context, our finding of a consistent association between protein background and HSA batch variability underscores the need for greater transparency in media composition, standardization of manufacturing processes, and stricter quality control for products containing human blood–derived components.

The development of fully chemically defined IVF media that exclude serum-derived components is becoming increasingly important for promoting reliable and standardized outcomes. Some manufacturers offer media supplemented with serum replacements; however, these formulations are generally less defined than HSA and are, therefore, more likely to introduce additional impurities. In our study, recombinant HSA (recHSA) did not introduce any of the analyzed contaminants, positioning it as a promising alternative. However, clinical use will require assurance of consistent purification and removal of residual host-cell proteins. Previous work by Bungum et al. ^48^ demonstrated that recHSA performs comparably to serum-derived HSA in embryo culture, yet it has not been widely adopted in clinical practice.

An important avenue for future research is the development of fully protein-free IVF media. Such synthetic media have already been tested in both animal ^49–52^ and human embryo culture ^53,54^, demonstrating their potential to support preimplantation development. These fully defined formulations would enable true standardization across IVF laboratories while also providing a highly controlled environment for studying the embryo secretome.

## CONCLUSIONS

This study provides compelling evidence that undeclared proteins are present in commercially available IVF culture media, even before embryo exposure. Through targeted and untargeted proteomic analyses, we demonstrate that these impurities contribute to significant lot-to-lot variability in media composition. Our results further show that the protein background is primarily introduced through supplementation with serum-derived human albumin of varying origin. These findings underscore the need for standardization in IVF media manufacturing and highlight the importance of including properly matched controls in proteomic studies of spent culture media.

## ACKNOWLEDGEMENTS

Funding for this research was provided by the Ministry of Health of the Czech Republic (NU22-08-00543), Grant Agency of Masaryk University (MUNI/A/1738/2024), and R&D funds of Reprofit International. The authors wish to thank the staff of Reprofit International for collecting media samples and gratefully acknowledge Dr. David Potěšil (Proteomics Core Facility, Central European Institute for Technology) for his help with the interpretation of untargeted proteomics data and critical review of the manuscript. We acknowledge the CEITEC Proteomics Core Facility of CIISB, Instruct-CZ Centre, supported by MEYS CR (LM2023042, e-INFRA CZ (ID:90254)). This work was supported from the European Union’s Horizon 2020 research and innovation program under grant agreement No 857560 (CETOCOEN Excellence). This publication reflects only the author’s view, and the European Commission is not responsible for any use that may be made of the information it contains. Authors thank the RECETOX Research Infrastructure (No LM2023069) financed by the MEYS CR for supportive background.

## DATA AVAILABILITY STATEMENT

The mass spectrometry proteomics data generated in this study have been deposited to the ProteomeXchange Consortium via the PRIDE partner repository with the dataset identifier PXD063245 (reviewer access token: bKwOy4H7oTKP). Data from the targeted proteomics experiment have been deposited in Panorama Public and are accessible via the permanent link: https://panoramaweb.org/GHihM3.url. This dataset is also available through ProteomeXchange under the identifier PXD063038, associated with the DOI: https://doi.org/10.6069/vkze-gj62. Reviewer access is available using the credentials: Password: FL!y?xOkG0IU+r

## AUTHOŔS ROLES

M.N.: Targeted proteomic analysis, data processing and interpretation, Manuscript and figure drafting; V.P.: Sample management, Conception of experiments, Data analysis and interpretation; T.B.: Statistical analysis and data interpretation; V.P.: Untargeted proteomics data analysis and interpretation; D.K., B.M: Sample collection and administration; S.K., P.O.: Critical reading of manuscript and publication approval; Z.H.: Project design, Data analysis, and interpretation, Manuscript writing. All authors discussed the results and commented on the final manuscript.

## DECLARATION OF GENERATIVE AI IN SCIENTIFIC WRITING

During the preparation of this work, the corresponding author used ChatGPT, an AI language model developed by OpenAI, in order to refine the clarity, coherence, and style of the manuscript text. After using this tool, the authors reviewed and edited the content as needed and take full responsibility for the content of the published article.

