## Supplementary Figures for "Human Serum Albumin as a Hidden Source of Variability in IVF Culture Media"

Running title: Protein contamination by serum-derived albumin

Markéta Nezvedová[1], Volodymyr Porokh[2,3], Tami Bočková[1,2], Václav Pustka[4], Drahomíra Kyjovská[2,3], Barbora Maierová[3], Soňa Kloudová[3], Pavel Otevřel[3], Zuzana Holubcová[2,3*]

[1] RECETOX, Faculty of Science, Masaryk University, 625 00 Brno, Czech Republic;

[2] Department of Histology and Embryology, Faculty of Medicine, Masaryk University, 625 00 Brno, Czech Republic;

[3] Reprofit International – Clinic of Reproductive Medicine and Gynecology, 603 00 Brno, Czech Republic

[4] Proteomics Core Facility, Central European Institute for Technology, Masaryk University, 625 00 Brno, Czech Republic;

* Corresponding author:, address: | Masaryk University, Faculty of Medicine, Department of Histology and Embryology, Masaryk University Campus, Kamenice 3, building F01B1, 62500 Brno, Czech Republic

### Supplementary Figure 1

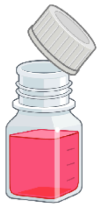

Irvine Scientific (FUJIFILM)

Continuous Single Culture Complete (CSCM-C)

Catalog ID: 90165

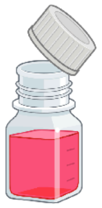

Cooper Surgical

SAGE 1-Step™

Catalog ID: 67010010

| Product | SCM samples | Commercial LOT number | BLANK samples |
|---|---|---|---|
| Irvine Scientific CSCM-C | | 011676 | → Lot 1 |
| Irvine Scientific CSCM-C | | 014943 | → Lot 2 |
| Irvine Scientific CSCM-C | | 015021 | → Lot 3 |
| Irvine Scientific CSCM-C | | 015052 | → Lot 4 |
| Cooper Surgical SAGE 1-Step™ | Lot #01 ← | 010781 | |
| Cooper Surgical SAGE 1-Step™ | Lot #02 ← | 011289 | |
| Cooper Surgical SAGE 1-Step™ | Lot #03 ← | 012759 | → Lot 5 |
| Cooper Surgical SAGE 1-Step™ | Lot #04 ← | 013580 | → Lot 6 |
| Cooper Surgical SAGE 1-Step™ | Lot #05 ← | 014694 | → Lot 7 |
| Cooper Surgical SAGE 1-Step™ | Lot #06 ← | 015751 | |
| Cooper Surgical SAGE 1-Step™ | Lot #07 ← | 017411 | → Lot 8 |
| Cooper Surgical SAGE 1-Step™ | Lot #08 ← | 019857 | |
| Cooper Surgical SAGE 1-Step™ | | 020554 | → Lot 9 |

**Supplementary Fig1: List of analyzed commercial IVF culture media.** Product information and lot mapping.

Supplementary Fig2

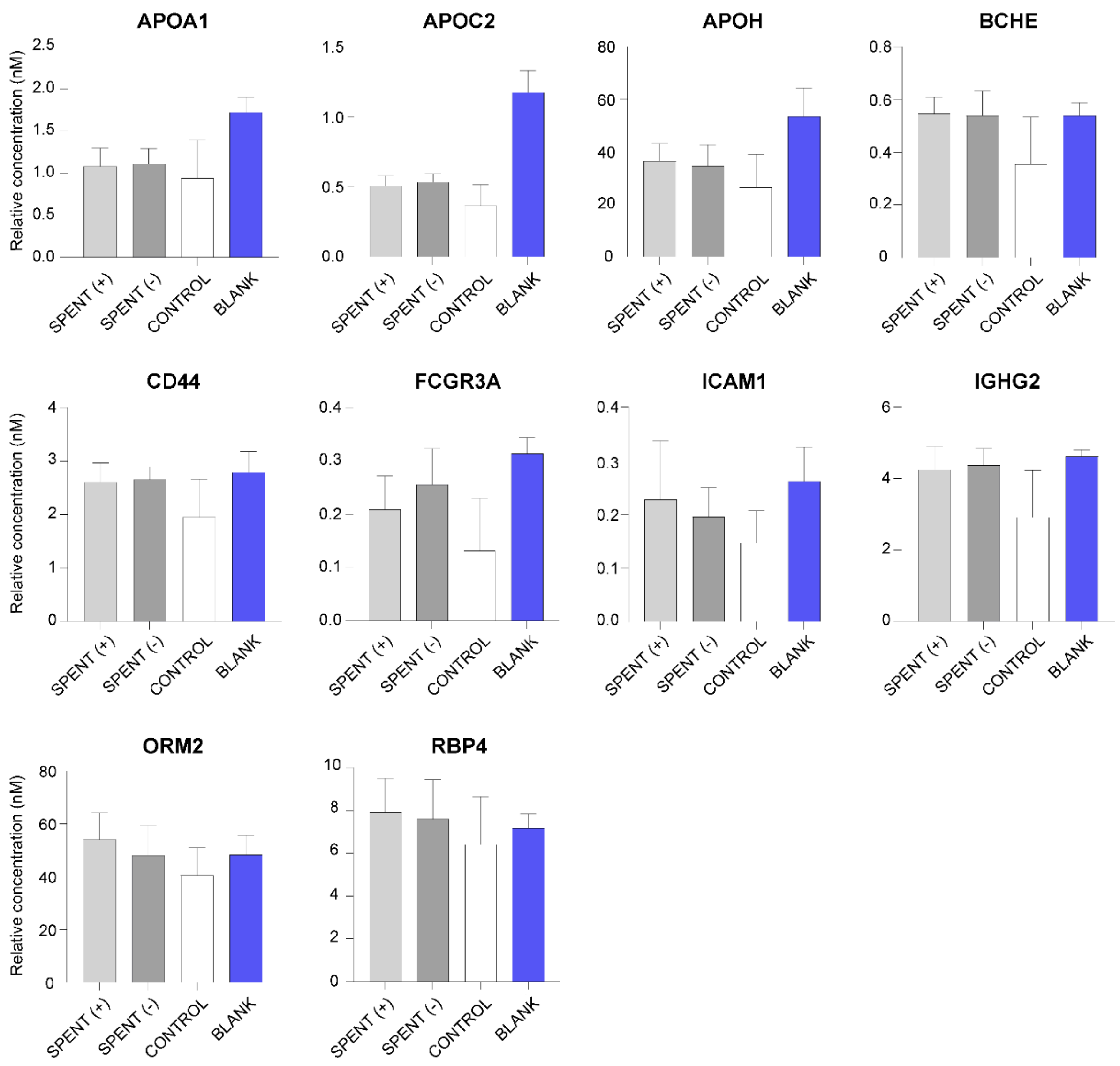


**Supplementary Fig2: Additional results from protein analysis of embryo culture media.** Results for 10 additional proteins detected by targeted proteomic analysis in a subset of SCM samples (n=17) from good-quality (SPENT+, n = 8) and poor-quality (SPENT−, n = 9) embryo cultures, corresponding controls (CONTROL, n = 5), and blank samples of unused media (BLANK, n = 8 technical replicates), complementing the findings presented in **Figure 2C**.

#### Supplementary Fig3

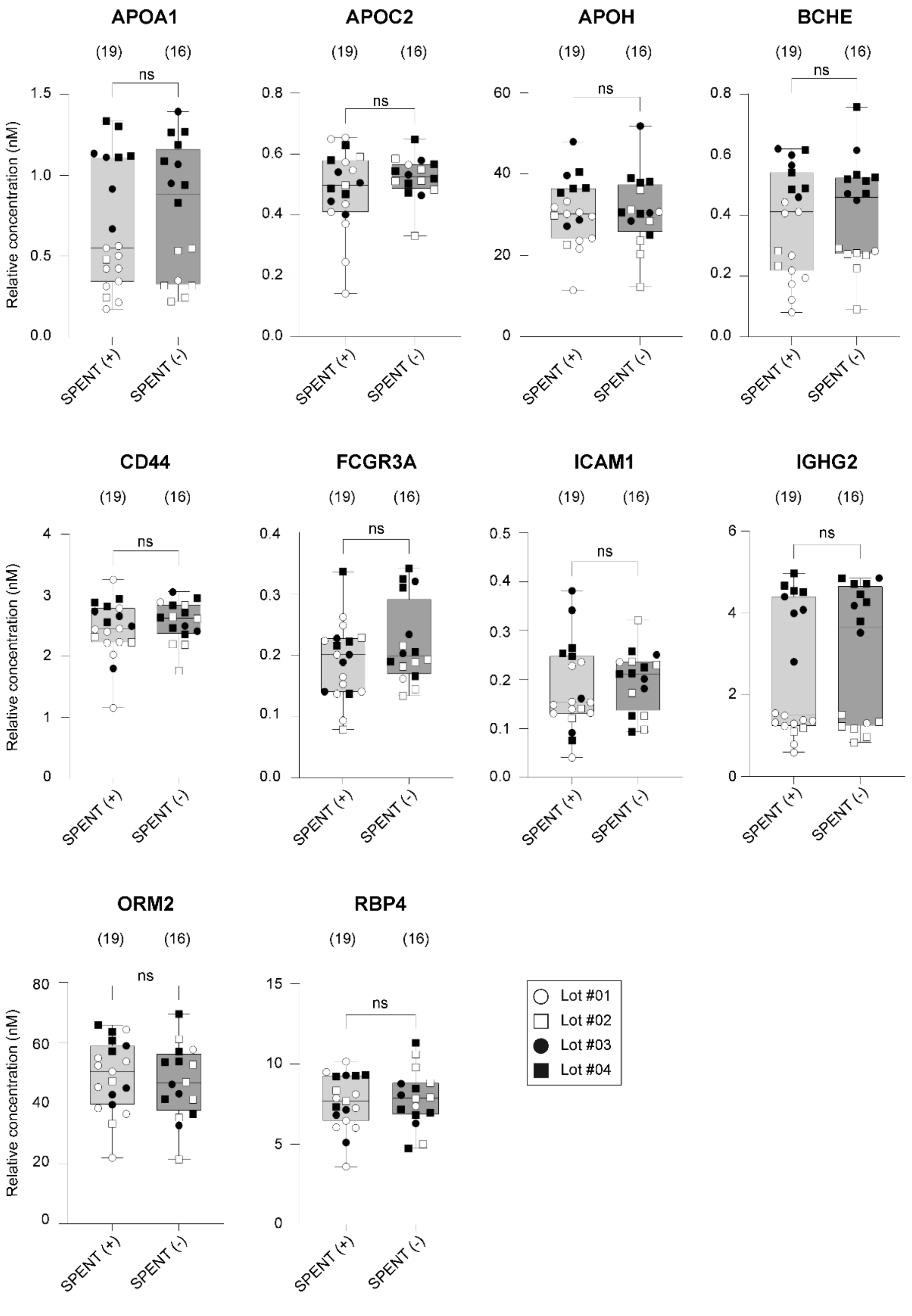


**Supplementary Fig3: Additional results from protein analysis of spent culture media with respect to the media Lot number.** Results from targeted protein analysis of additional 10 proteins detected in SCM from good-quality (SPENT+) and poor-quality (SPENT−) embryos, categorized based on the Lot number (Lot #01-04) of the culture media used. "ns" indicates statistically non-significant differences (Mann-Whitney test, $p>0.05$). (n) denotes the number of samples per group. These data complement those presented in **Fig3A**.

### Supplementary Fig4

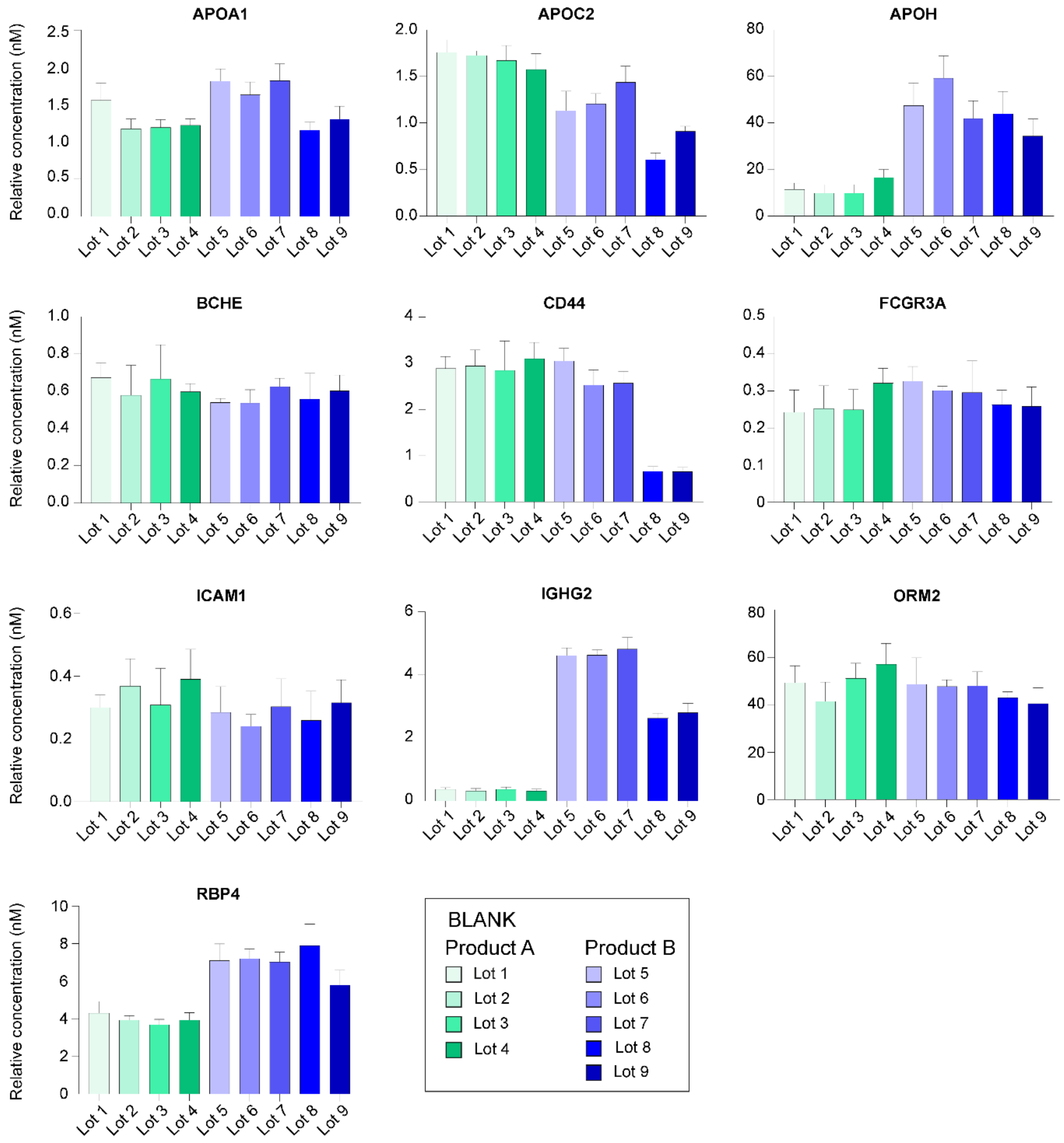


**Supplementary Fig4: Additional results from protein analysis of blank media samples.** Results for the additional 10 proteins detected by targeted analysis in 9 Lots of embryo culture media from two different producers—Product A (shades of green) and Product B (shades of blue). These data complement those presented in **Fig4A**.

### Supplementary Fig5

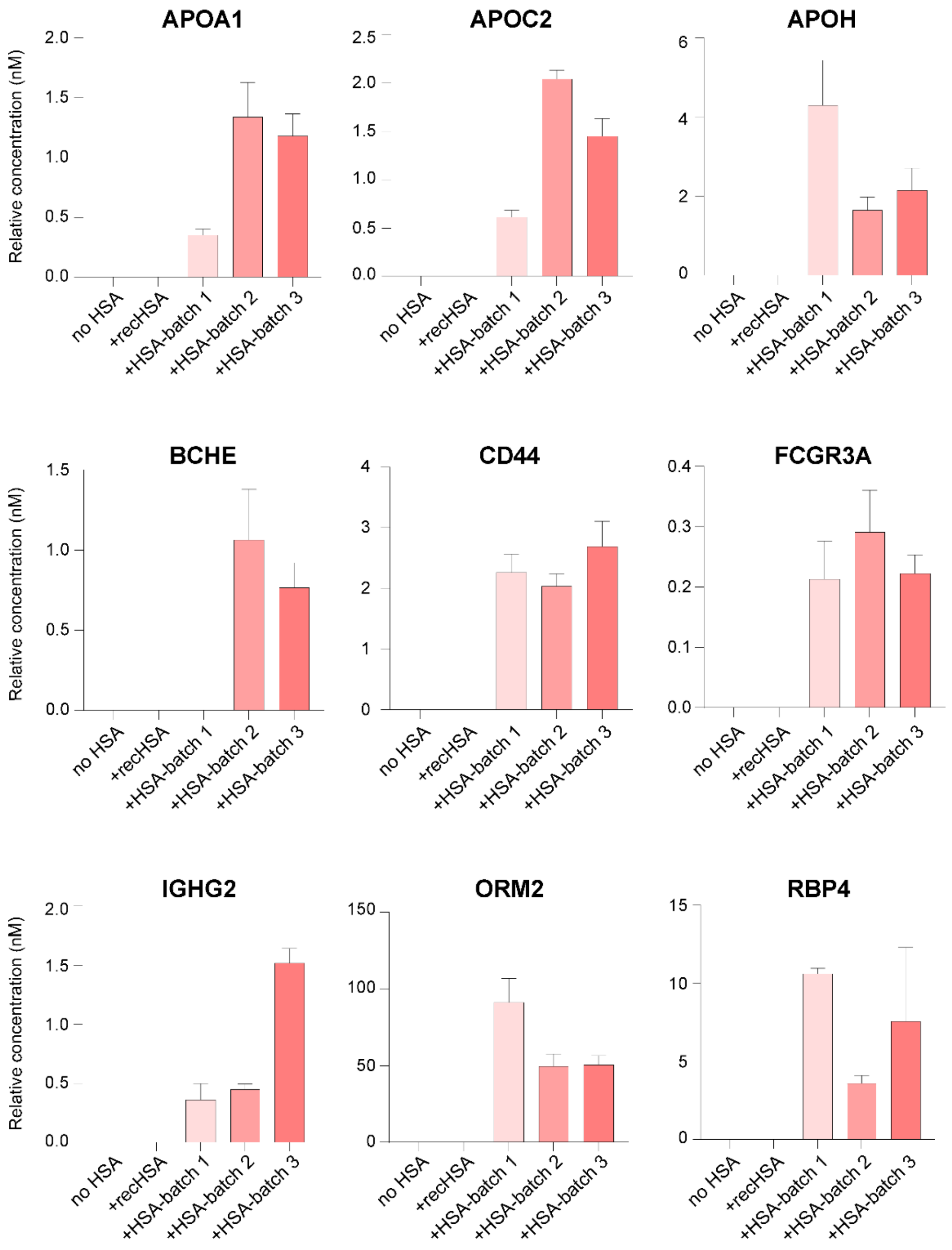


**Supplementary Fig5: Additional results from the investigation of protein background sources in embryo culture media.** Results for the additional 9 analyzed proteins detected by targeted analysis in all types of samples from the given experiment. These data complement those presented in **Fig5C**.

### Supplementary Fig6

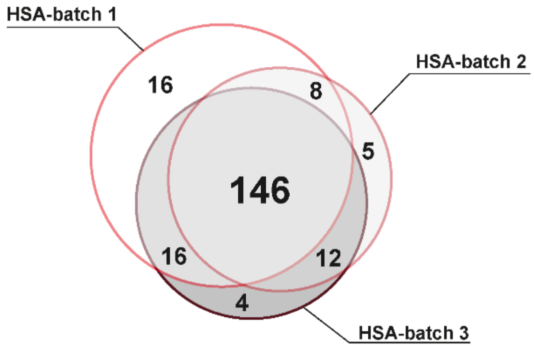


**Supplementary Fig6: Protein composition of IVF media supplemented with serum-derived HSA.** Untargeted analysis of protein-free media supplemented with three different batches of serum-derived HSA (batches 1–3) revealed that the majority of proteins were shared across groups (central intersection), while some protein markers were unique to individual HSA batches.
